# Effect sizes of somatic mutations in cancer

**DOI:** 10.1101/229724

**Authors:** Vincent L. Cannataro, Stephen G. Gaffney, Jeffrey P. Townsend

## Abstract

A major goal of cancer biology is determination of the relative importance of the genomic alterations that confer selective advantage to cancer cells. Tumor sequence surveys have frequently ranked the importance of substitutions to cancer growth by *P* value or a false-discovery conversion thereof. However, *P* values are thresholds for belief, not metrics of effect. Their frequent misuse as metrics of effect has often been vociferously decried. Here, we estimate the effect sizes of all recurrent single nucleotide variants in 23 cancer types, quantifying relative importance within and between driver genes. Some of the variants with the highest effect size, such as EGFR L858R in lung adenocarcinoma and BRAF V600E in primary skin cutaneous melanoma, have yielded remarkable therapeutic responses. Quantification of cancer effect sizes has immediate importance to the prioritization of clinical decision-making by tumor boards, selection and design of clinical trials, pharmacological targeting, and basic research prioritization.

## STATEMENT OF SIGNIFICANCE

We calculated the effect size for all recurrent single-nucleotide variants in 23 cancer types, quantifying the relative importance of the mutations driving cancer cell replication and survival. This quantification provides a means to prioritize basic research, inform decisions by tumor boards, and inform design of clinical trials.

## INTRODUCTION

Since the advent of whole-exome and whole-genome sequencing of tumor tissues, studies of somatic mutations have revealed the underlying genetic architecture of cancer (1), producing ordered lists of significantly mutated genes whose ordering implies their relative importance to tumorigenesis and cancer development. Typically, differentiation of selected mutations from neutral mutations is performed by quantifying the over-representation of mutations within specific genes in tumor tissue relative to normal tissue, and disproportionate prevalence of somatic mutations in a gene has been taken as *prima facie* evidence of a causative role for that gene. Two quantifications have implicitly ordered the importance of discovered cancer “driver” genes: the prevalence of the mutation among tumor tissues sequenced from that tumor type (2,3), the statistical significance (*P* value) of the disproportionality of mutation frequency (4), or both (1). Versions of these metrics have shifted from simple ranks by mutation prevalence in a tumor population (5–7) to calculation of statistical significance of mutation prevalence over genome-wide context-specific background mutation rates (8–10), to ratios of the prevalence of nonsynonymous and synonymous mutations (11), to *P* values based on a gene-specific mutation rate and a diversity of genomic data (12). Although the approaches used to calculate *P* values have become more sophisticated, *P* values are not an appropriate metric for quantifying the vital role of genes or their mutations to tumorigenesis and cancer development, as *P* values are thresholds for belief (13) and not metrics of effect (14,15). Failure to report effect sizes is a persistent issues in the scientific and biomedical literature that can massively misdirect research and health care decision-making (16–20).

While prevalence of a somatic mutation in a cancer type has important consequences for biomarker studies (21) and identification of therapeutic population for a targeted therapy (22), there is only a correlative—rather than causal—link between prevalence of mutation and its contribution to tumorigenesis and cancer development. The lack of causal linkage is easily seen by considering the mutated genes that, in spite of their high prevalence in tumor populations, are universally regarded as false positives. For example, the gene *TTN* is a structural protein of striated muscle. Because it is long, because it replicates relatively late in the synthesis phase of mitosis, and because it is inaccessible to transcription-mediated repair in non-muscle tissues, it has a high mutation rate and is frequently mutated in non-sarcoma cancer tissues. While *TTN* is an extreme example—sometimes showing up at the top of lists ordered by prevalence of genic mutation (11,12)—it exemplifies the problem with using mutation prevalence as a proxy for importance. Any consideration of whether mutated genes are contributing to tumorigenesis and cancer development—or of the degree to which they are contributing—must address the issue of their underlying mutation rate.

The appearance of such biologically implausible genes in ranked mutation lists prompted the development of increasingly sophisticated statistical approaches designed to “weed out” false-positives via calculation of a *P* value that accounted for gene length and background mutation rate. The classical evolutionary biology approach is to use the frequency of synonymous site mutations in each gene as a proxy for mutation rate. As in the divergence of species, synonymous site mutations are presumably neutral (or nearly so) to the success of cancer lineages during the divergence from normal to resectable tumor. As the number of mutations observed in a given gene is typically much smaller in the somatic evolution of cancer than is observed in the divergence between most species, use of synonymous sites within a single gene leads to many genes with zero synonymous mutations in most cancers, and an ineffective calculation of *P* value. Alternative approaches currently in use obtain a reasonably robust estimate of genic mutation rate using correlates such as gene expression levels, chromatin states, and replication timing, and are largely successful at excluding known false positives (12).

In genomic tumor surveys, the sample size of tumors varies among studies, posing a problem for comparison of *P* values within or between cancer types (14,15). An even more serious issue with using *P* values for ranking genes or mutations arises from the same source that obviates use of genic mutation prevalence: the confounding effect of mutation rate. Because the “sample size” of overall mutations in a gene is dictated by the genic mutation rate, it is much easier for genes with high mutation rates not only to reach high genic prevalence, but also to reach statistical significance despite small effect sizes. While approaches accounting for genic mutation rates will eliminate false positives (12), and the *P* value will serve to exclude genes like *TTN* that have no role in tumorigenesis and cancer development, the rank order by *P* value of genes that do have a role in tumorigenesis and cancer development will remain highly affected by mutation rate. Genes with higher mutation rate will (correctly) be more likely to achieve statistical significance, and thus will appear deceptively high on a ranked list whose ordering suggests importance in tumorigenesis and cancer development.

Because genic mutation prevalence and *P* value inadequately capture importance to tumorigenesis and cancer development, another metric must be appropriate. To provide an evaluation of the relative importance of mutations in diverse cancer types to tumorigenesis and cancer development, we called on an understanding of the development of cancer as an evolutionary process (23,24), and adapted some straightforward insights from classical evolutionary theory. The cognate metric in evolutionary theory for quantifying importance to tumorigenesis and cancer development is the selective effect of the mutation on the cancer lineage. The appropriateness of this metric is fairly easy to recognize. While mutations are the ultimate source of variation contributing to tumorigenesis, we do not conduct genomic tumor sequence surveys to discover neutral mutation rates. We conduct them to determine which mutations spread within cancer tissues because of the effects of mutations on proliferation and survival. Mutation rate is a confounding phenomenon: when it is high, it also increases prevalence of mutations. Because silent site substitutions and other correlates of baseline mutation rate provide a means to independently differentiate silent mutation rate from the impact of natural selection within the tumor, selection intensities can be estimated, providing the effect sizes of each mutation.

## RESULTS

We calculated cancer effect sizes by comparing the rate of observed substitutions to the rate that substitutions would be expected to arise in the absence of selection (25). In accordance with population genetic theory, we specify that the rate neutral mutations arise and the rate that they fix as substitutions within tumors are equivalent, and that non-neutral mutations arise at a consistent rate. Thus any increase in the flux of substitutions among tumors of a particular context above the baseline silent rate would be the appropriate estimate of the intensity of selection on that mutation within the tumor population (Methods). These selection intensities quantify the survival and proliferative advantage conferred by variants, facilitating comparisons of the relative importance of drivers among and within tumor types. For instance, even though lung tissues encounter similar mutagens and hence mutations arise through similar mutational signatures, LUAD and LUSC variants conferring the largest effect sizes are markedly different among the two cancers (Fig. 1). Moreover, within a tumor type, the selection intensity decouples the effects of mutation rate on frequency: even though KRAS G12C variants are over twice as prevalent as EGFR L858R variants in LUAD tumors, EGFR L858R is estimated to have a higher effect size, due to its much lower baseline mutation rate. Furthermore, the relative selection intensity among variants within a single gene reveals whether ‘hotspots’ of somatic variation are purely driven by mutational processes or rather under differential (or similar) selective pressure. For instance, the V157F, R158L, R175G, G245C, and E298* variants in TP53 in LUSC tumors exhibit markedly different prevalences attributable to different trinucleotide mutation contexts, yet resulting in remarkably similar estimated effect sizes.

**Figure 1:**
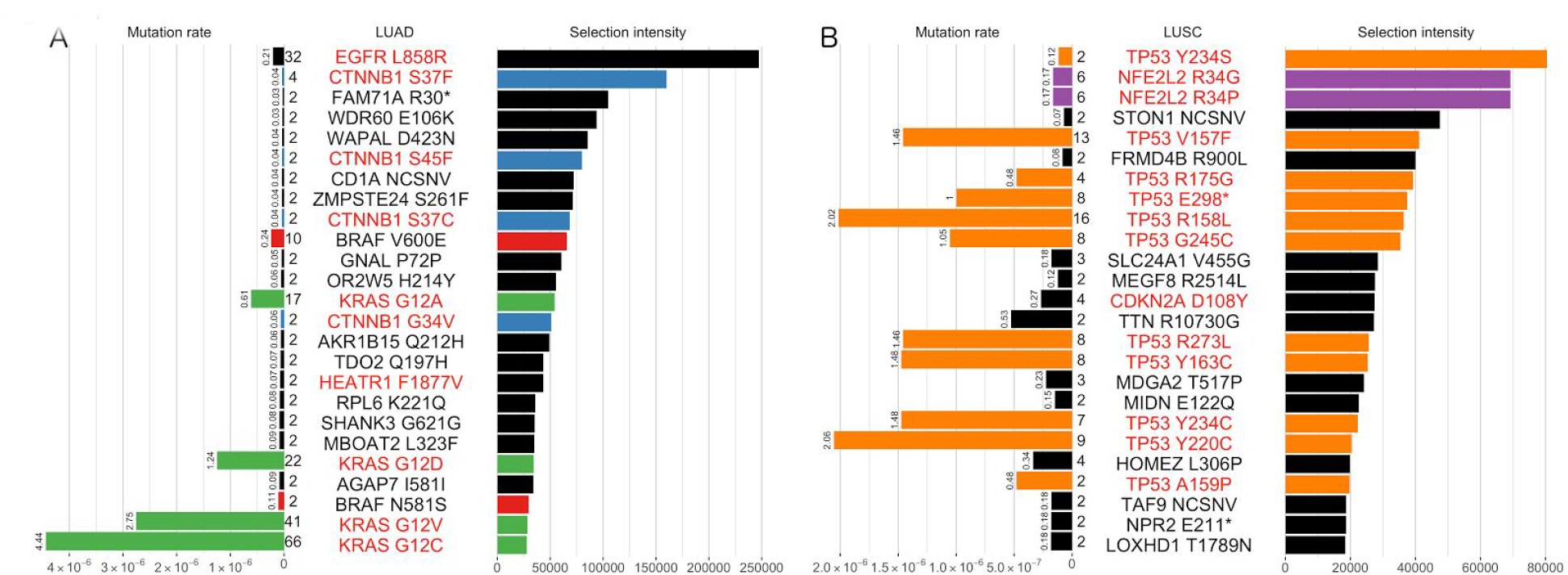
The 25 single-nucleotide variants with the highest selection intensity in LUAD and LUSC tumors, ranked by selection intensity, which corresponds to the length of the bars to the right of the variant names. Length of bars to the left of variant names corresponds to mutation rate, and mutation rate (× 10^−6^) is listed at the top of the bars. Prevalence of variants among 675 LUAD and 600 LUSC tumors are listed to the left of the variants. Red variant labels refer to genes deemed significantly mutated by `MutSigCV` (*Q* < 0.1) and variants occurring in genes present twice in the top 25 have uniquely colored bars. Asterisks denote nonsense mutations.

This calculation yielded cancer effect sizes for all fixed substitutions (Methods, Figure S1, Table S1) that quantify contribution to the cancer phenotype within 23 tumor types. Their relative rank corresponds to their relative importance within the respective tumor types. Several common known oncogenic substitutions, such as BRAF V600E in SKCMP and EGFR L858R in LUAD (26,27), and substitutions in known tumor suppressor genes, such as APC in READ and TP53 in HPV-negative HNSCC, are highly selected, and those genes are also typically determined as significantly mutated (e.g., by MutSigCV(12)). However, genes determined to be statistically significantly altered in cancer are also well-dispersed within a large range of site-specific cancer effect sizes (Fig. 2), illustrating how discrepant gene-specific *Q* values are with site-specific cancer effect sizes. Several substitutions within genes that are not statistically significantly over-mutated are interspersed among more prevalent substitutions within genes that are estimated to be significantly mutated, for instance Mastermind-like3 (MAML3) G1069A in READ, a protein that binds to and stabilizes the DNA-binding complex of the Notch intracellular domain (28) and Nuclear factor (erythroid derived 2)-like 2 (NFE2L2) R34G in UCEC, a protein that is believed to play a causative role in squamous cell lung cancer (29,30). Indeed, substitutions in this gene comprise two of the top three most selected substitutions within our analysis of lung squamous cell carcinoma (Fig. 2).

**Figure 2:**
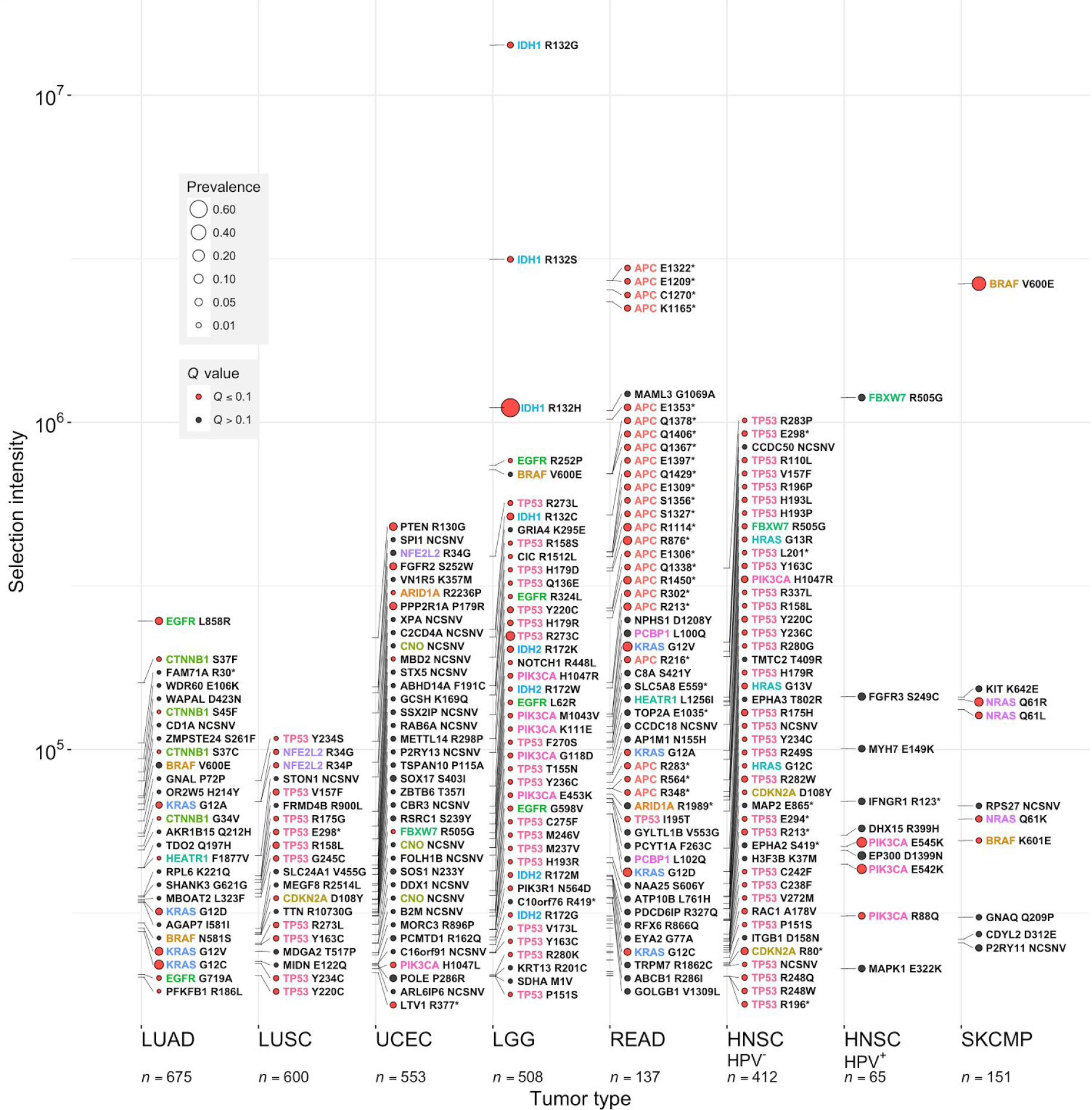
Cancer effect sizes of recurrent somatic substitutions in eight of the 23 cancer types analyzed. Effect sizes greater than 1 × 10^3^ are indicated by ticks along the tumor-type axes. The highest 50 effect sizes are labeled within each tumor. Names of genes that have more than one mutation within or between tumors are uniquely colored. Genes deemed significantly burdened with mutation (12) are depicted by a red circle next to mutation labels, and the prevalence of each substitution is represented by the size of this circle. LUAD: Lung adenocarcinoma; LUSC: Lung squamous cell carcinoma. UCEC: Uterine corpus endometrial carcinoma; LGG: Brain Lower Grade Glioma; HNSC: Head and neck squamous cell carcinoma, broken into HPV positive and HPV negative tumor samples (methods); SKCMP: Skin cutaneous melanoma (primary); READ: Rectum adenocarcinoma. NCSNV refers to a non-coding single nucleotide variant outside an exon (e.g. 5’ or 3’ UTRs). HPV^+^ and HPV^-^ HNSCC have been demonstrated to have significantly different genetic architectures (43); thus, they are presented separately. 108 LUAD, 108 LUSC, 23 UCEC, and 47 SKCMP tumors from the Yale-Gilead collaboration are included in the plot.

## DISCUSSION

Here, we have argued that frequencies of mutation and *P* values are not an appropriate metric for the importance of somatic variants in tumor growth. We derive the appropriate metric—the cancer effect size of mutations. This effect size, quantifying the intensity of selection on mutations in cancer cells in patients, conveys the replicative and survival benefit conferred by genetic variants, and therefore is a direct metric of the contribution of a variant to the cancer phenotype. Using our approach, we estimated the effect size of all recurrent SNVs in 23 cancer types, re-evaluating their importance across SNV effect size to cancer in disparate tumor types projected to account for approximately 82% of all new cancer cases within the United States in 2017 (31).

Current approaches using conservative *P* values are particularly underpowered to detect genes that are of high importance to tumorigenesis and cancer development in some cancer cases because the site or sites conveying the relevant phenotype are mutated at low rates. For instance, FBXW7 R505G was estimated to have the highest selection intensity in HPV^+^ HNSCC, and BRAF V600E was estimated to have the fifth highest selection intensity in LGG and the tenth highest selection intensity in LUAD, but both of these genes were classified as not significantly mutated at the gene-level within these three cancer types. Mutations within these two well-known oncogenes were estimated to confer large effect sizes, and these genes were determined to be significantly mutated in other cancer types, yet their importance in patients who have these rarer somatic variants within HPV^+^ HNSCC, LGG, and LUAD has been poorly illuminated by gene-wide analyses of statistically significant mutation burden across patients. Thus, calculating the cancer effect size at the level single nucleotide variants identifies drivers that have low prevalence but large effect in different tumor types.

Identification of these low-prevalence high-effect drivers is increasingly important as precision targeting of therapeutics becomes increasingly integrated into clinical trial design. Targeted therapies are often developed for, and necessarily tested in, a single tumor type with the targeted variant at high frequency. However, the same variant often exists at lower prevalence in other tumor types. Quantifications of the cancer effect size can guide the selection and design of clinical trials to target small populations that can benefit from targeted therapeutics developed for other cancer types. Targets for therapies that were originally developed for variants at high prevalence in one tumor type are expected to be effective when treating the same variant in a secondary tumor type if the variant has a similarly high selection intensity in the second tumor type, even if it is at low prevalence in the newly considered tumor type.

The effect sizes of somatic mutations in cancer should play key roles in clinical decision-making, providing crucial insight into the relative upper limits of the efficacy of precision-targeted treatments. When a therapy abrogates novel oncogenic function primarily by disabling a gain-of-function mutation, the upper limit of the efficacy of the precision-targeted treatment will be dictated by the selection intensity that somatic variant imparted to the cancer cell lineage. Therefore, while incorporating other knowledge regarding the pharmacokinetics, efficacy, and side effects, and the evolution of resistance to therapies, precision treatments can be selected in clinical decision-making on tumor boards to target mutations with the greatest cancer effect size. That is, the effectiveness of a targeted therapy is expected to scale with the selection intensity of the target for any therapy that specifically inhibits the selective advantage conferred by the mutant allele. Similarly, cancer effect size can be used in the same manner to prioritize targeted development of new therapeutics, indicating the upper limit of effect for a perfect therapeutic ameliorating an oncogenic mutation.

The relative effect sizes of cancer mutations can inform nearly every aspect of basic research related to oncology, including which genes and pathways deserve greater attention in basic research. While network interactions, epigenetic and tumor microenvironment interactions, and aspects of cellular differentiation and cellular plasticity mean that quantification of the selective effect of mutations does not provide an upper bound on the importance of a gene or pathway in the molecular biology of cancer, its quantification does provide a lower bound, as the potential role of genes with a somatic variant can be no lower than the selection intensity the variant imparts. It does not escape our notice that here we only calculate the effect size of single nucleotide variants. Importantly, effect sizes of other mutational processes, such as copy number alterations or epigenetic modifications, could be similarly calculated once estimates of intrinsic mutation rate in the absence of selection are identifiable for these processes.

## METHODS

### Data acquisition and processing

Data were either obtained either from The Cancer Genome Atlas (TCGA) projects downloaded from the National Cancer Institute’s Genomic Data Commons (32), or derived from whole exome sequencing of tumors as part of collaboration with Gilead Sciences (Table S2). All TCGA data used in this analysis were from GDC version gdc-1.0.0 and relevant UUID are found in Table S3.

All TCGA data were first converted to hg19 coordinates using the `liftOver` function of the `rtracklayer` R package(33). To obtain consistency between gene symbols used in `MutSigCV` and in mutation data, symbols in the `MutSigCV` default covariates files were mapped to HUGO symbols. Non-HUGO symbols were mapped using substitutions from the NCBI Gene database using unambiguous ‘synonym’ matches; unambiguous ‘previous symbol’ matches; and manual lookups. CDS coordinates for each gene were obtained from UCSC’s hg19 annotation database. The `MutSigCV` covariates files, and gene annotation files used in our analyses can be found at https://github.com/Townsend-Lab-Yale/SNV_selection_intensity. Nucleotide variants one or two positions apart in the same tumor sample were removed from the analysis to ensure that we analyzed only single nucleotide variants. Head and neck squamous cell tumors were designated as HPV positive if they both contained greater than 100 HPV RNA viral transcript reads per hundred million (RPHM) (34) and were designated positive in clinical data obtained from The Broad GDAC Firehose (35), and tumors were designated HPV negative if they contained less than 100 HPV RNA viral transcript RPHM and were designated negative in clinical data obtained from The Broad GDAC Firehose.

### Calculating mutation rate and selection intensity

We define the selection intensity of a single nucleotide variant as the ratio of the flux of fixed mutations to the expected flux of fixed mutations in the absence of selection. We estimate the expected rate of fixed mutations in the absence of selection by using `MutSigCV` to calculate the silent mutation rate for each gene. We use the median gene expression from cell lines derived from each analyzed tissue type (Table S3) as the `MutSigCV` expression covariate for that tissue type. We then normalized this gene-level rate at every site for every mutation across each gene given the specific trinucleotide mutation profile of the tissue, such that the average rate of mutations among all trinucleotide combinations in each gene is equal to the average rate calculated by `MutSigCV`. The trinucleotide profile of a tissue is calculated as the average of trinucleotide COSMIC signatures among non-recurrent mutations calculated by the `deconstructSigs` R package (36) for all tumors with over 50 single nucleotide variants.

Formally, we define the rate of non-neutral substitutions, λ, as *N*μ × *u*(*s*), where *N* is the effective population size of the tumor, μ is the mutation rate of a substitution, and *u*(*s*) is the probability the mutation fixes within the population, which is a function of the selection coefficient of the mutation, *s*. The probability that a silent mutation fixes is the inverse of the population size within which it fixes (37), and thus rate of non-neutral substitutions, divided by the rate of neutral substitutions, is 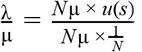 or *N* × *u*(*s*), the selection intensity and effect size of the mutation. The term selection intensity is used as a direct parallel to the classic derivation of “scaled selection coefficient” or “selection intensity” in the population genetics literature (38–41). Because a tumor sequence is a single snapshot in time, we can only detect one substitution event per site per tumor. To correct for this issue of detection, we define the rate of substitution, λ, as the Poisson rate of occurrence of one *or more* observed fixation events, i.e. the value of λ that maximizes 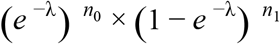, where *n*_0_ is the number of tumors without any substitution in that gene and *n*_1_ is the number of tumors with the specified substitution at that site. We define *n*_0_ as the number of tumors without any substitutions in that gene, instead of in that gene at that specific site, because doing so accounts for the reduced or eliminated selective pressure on additional mutation for that gene when other sites are already mutated. We define substitutions as single nucleotide variants that are observed in tumors from more than one patient (recurrent) within our dataset, and we calculate the selection intensity of the recurrently mutated substitutions to minimize the probability of analyzing passenger mutations.

Scripts used to perform this analysis are available online at https://github.com/Townsend-Lab-Yale/SNV_selection_intensity

## Supplementary Information

available online.

## Acknowledgements

This project was supported by Gilead Sciences, Inc. We also thank the Yale High Performance Computing Center for providing computational resources.

## Author contributions

VLC and JPT designed the analyses. VLC performed the analyses with assistance from SGG. JPT, VLC, and SGG wrote the manuscript.

## TABLES

Tables S1, S2, and S3 available as a supplement to the manuscript.

## TABLE LEGENDS

Table S1: Tab delimited text file containing the selection intensity of all recurrent single nucleotide variants in our analysis. Informative columns also include the amino acid position of the variant (AA_pos), the amino acid reference (AA_Ref), the amino acid change (AA_Change), and prevalence of the variant among tumors (freq), the silent substitution rate of the variant (mu), and the `MutSigCV` *P* and *Q* values for that gene (MutSigCV_p and MutSigCV_q). When the variant is outside of an exon, the position is given in terms of the nucleotide position on the chromosome (Nucleotide_position), and the reference is the nucleotide reference (Nuc_Ref) and the change is the nucleotide change (Nuc_Change).

Table S2: Exome sequencing data from the Yale-Gilead collaboration displayed in accordance with Mutation Annotation Format (MAF) Specifications (42) in a tab-delimited text file.

Table S3: Metadata on the TCGA data used in this analysis in a tab-delimited text file. Includes NCI UUID for all files, and the gene expression tissue files used for each tumor type.

## FIGURE LEGENDS

**Figure S1:**
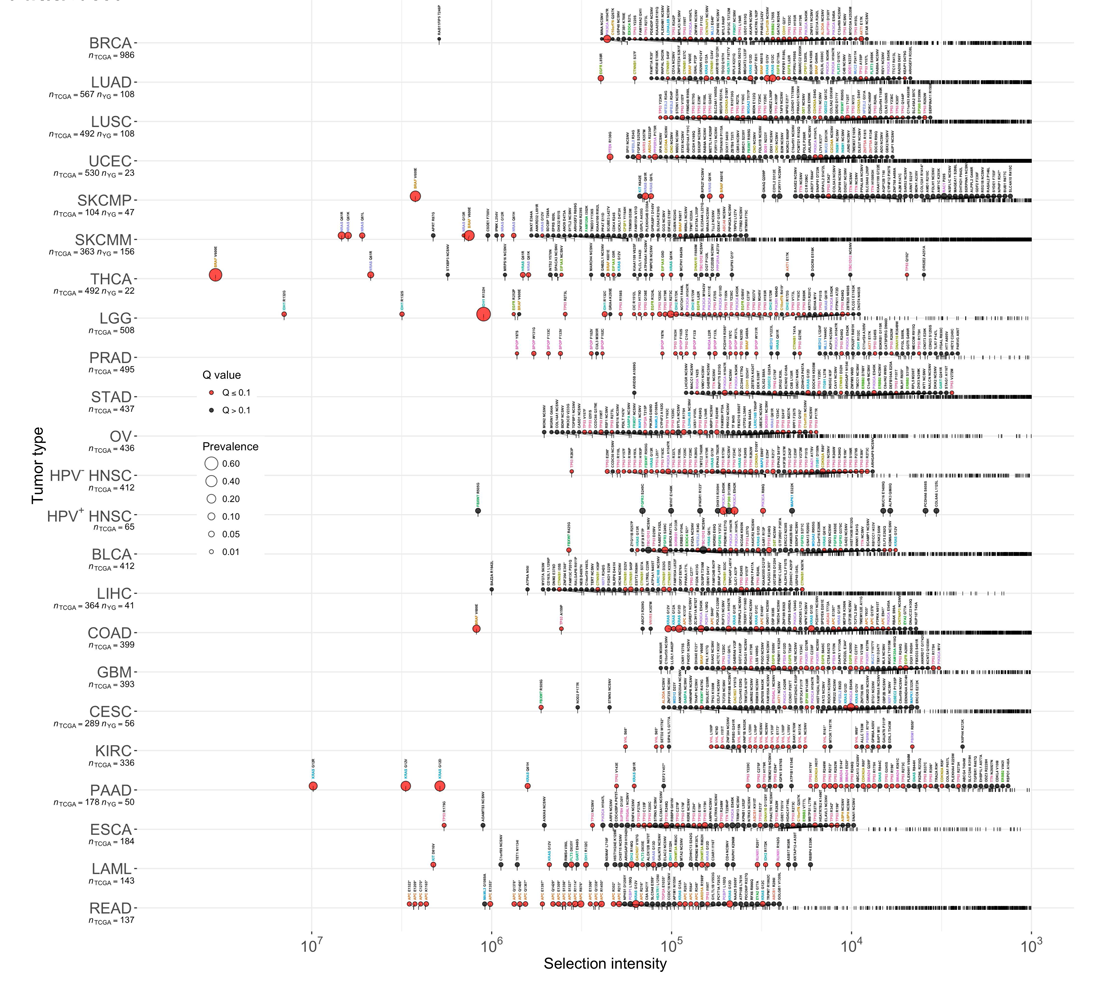
Selection intensities of recurrent somatic substitutions in 23 cancers. BLCA: Bladder Urothelial Carcinoma; BRCA: Breast invasive carcinoma; Cervical squamous cell carcinoma and endocervical adenocarcinoma; COAD: Colon adenocarcinoma; ESCA: Esophageal carcinoma; GBM: Glioblastoma multiforme; HNSC: Head and neck squamous cell carcinoma, broken into HPV^+^ and HPV^−^ tumor samples using criteria described within the Methods section; KIRC: Kidney renal clear cell carcinoma; LAML: Acute Myeloid Leukemia; LGG: Brain Lower Grade Glioma; LIHC: Liver hepatocellular carcinoma; LUAD: Lung adenocarcinoma; LUSC: Lung squamous cell carcinoma; OV: Ovarian serous cystadenocarcinoma; PAAD: Pancreatic adenocarcinoma; PRAD: Prostate adenocarcinoma; READ: Rectum adenocarcinoma; SKCM: Skin Curaneous Melanoma, broken into primary skin tumors and metastatic skin tumors; STAD: Stomach adenocarcinoma; THCA: Thyroid carcinoma; UCEC: Uterine corpus endometrial carcinoma. 108 LUAD, 108 LUSC, 23 UCEC, 47 SKCMP, 156 SKCMM, 22 THCA, 41 LIHC, 56 CESC, and 50 PAAD tumors from the Yale-Gilead collaboration are included in the plot.

